# Cigarette smoke exposure and inflammatory signaling increase the expression of the SARS-CoV-2 receptor ACE2 in the respiratory tract

**DOI:** 10.1101/2020.03.28.013672

**Authors:** Joan C. Smith, Erin L. Sausville, Vishruth Girish, Monet Lou Yuan, Kristen M. John, Jason M. Sheltzer

## Abstract

The factors mediating fatal SARS-CoV-2 infections are poorly understood. Here, we show that cigarette smoke causes a dose-dependent upregulation of Angiotensin Converting Enzyme 2 (ACE2), the SARS-CoV-2 receptor, in rodent and human lungs. Using single-cell sequencing data, we demonstrate that ACE2 is expressed in a subset of secretory cells in the respiratory tract. Chronic smoke exposure triggers the expansion of this cell population and a concomitant increase in ACE2 expression. In contrast, quitting smoking decreases the abundance of these secretory cells and reduces ACE2 levels. Finally, we demonstrate that ACE2 expression is responsive to inflammatory signaling and can be upregulated by viral infections or interferon treatment. Taken together, these results may partially explain why smokers are particularly susceptible to severe SARS-CoV-2 infections. Furthermore, our work identifies ACE2 as an interferon-stimulated gene in lung cells, suggesting that SARS-CoV-2 infections could create positive-feedback loops that increase ACE2 levels and facilitate viral dissemination.

## Introduction

In December 2019, a novel respiratory disease emerged in a seafood market in Wuhan, China^1^. Genomic sequencing demonstrated that the causative agent was a highly-contagious coronavirus, since named SARS-CoV-2^2,3^. The disease, called COVID-19, rapidly spread worldwide, and as of April 2020, more than 3 million people have been infected and more than 200,000 people have died^4^. No clinically-validated treatment or vaccine for COVID-19 is currently available. Thus, understanding the factors that mediate susceptibility to SARS-CoV-2 is crucial for controlling disease transmission.

Molecular analysis has begun to shed light on how SARS-CoV-2 infections occur. Like a related coronavirus that emerged in 2003^5^, SARS-CoV-2 enters human cells by binding to the extracellular domain of Angiotensin Converting Enzyme 2 (ACE2)^3,6^. Importantly, ACE2 is both necessary and sufficient for infection by SARS-CoV-2: ACE2-targeting antibodies block viral uptake in permissive cells while transgenic expression of human ACE2 allows viral entry in nonhuman cells. ACE2 normally functions in the renin-angiotensin system (RAS) by cleaving the vasoconstrictive hormone angiotensin-II into the vasodilator angiotensin 1-7^7^. Sequestration of ACE2 by coronavirus dysregulates the RAS pathway, contributing to morbidity^8^. Additionally, ACE2 levels are capable of influencing disease progression: among a cohort of mice engineered to express human ACE2, mice expressing the highest levels of ACE2 mRNA exhibited the shortest survival time following coronavirus exposure^9^. Thus, the regulation of ACE2 expression likely has a significant effect on SARS-CoV-2 susceptibility.

Epidemiological studies have identified multiple demographic features that correlate with the severity of clinical COVID-19 cases. While fewer than 5% of SARS-CoV-2 infections are fatal^10,11^, men and elderly patients are particularly at risk of developing severe disease^12–15^. Additionally, cigarette smokers with COVID-19 are significantly more likely to develop critical illnesses that require aggressive medical intervention^16,17^. For instance, in a study of 1,099 patients with laboratory-confirmed COVID-19, 12.3% of current smokers required mechanical ventilation, were admitted to an ICU, or died, compared to only 4.7% of non-smokers^13^. The causes underlying these differences in outcome are at present unknown.

Several clinical features have also been identified that can differentiate between patients with severe and non-severe coronavirus infections. Notably, a dysregulated immune response has been identified as a crucial mediator of COVID-19 mortality^18^. Patients who present with elevated levels of inflammatory cytokines are more likely to develop critical illnesses^19–21^. These so-called “cytokine storms” cause an increase in vascular permeability that facilitates immune cell efflux into affected tissues, but may also worsen pneumonia^22^. How this immunopathology affects the regulation of the host factors required for coronavirus infections is poorly understood.

## Results

### ACE2 levels in mammalian lungs are not strongly affected by age or sex

In order to study factors that could potentially influence susceptibility to SARS-CoV-2 infection, we investigated the expression of the coronavirus receptor ACE2. We first assessed the expression of ACE2 in a variety of rodent and human tissues (Figure S1 and Table S1)^23–26^. ACE2 was expressed at high levels in mouse, rat, and human kidneys, consistent with its role as a regulator of the RAS pathway. ACE2 was also expressed in rodent and human lungs, the predominant location of coronavirus infections^27^. Interestingly, significant ACE2 expression was also evident in the mouse and human small intestine. Viral RNA has been detected in stool samples from patients with COVID-19^28^, and gastrointestinal symptoms have been reported in a subset of affected individuals^13^, suggesting a potential alternate route for SARS-CoV-2 transmission. However, as SARS-CoV-2 appears to be primarily spread through viral inhalation^27^, we focused our study on factors that affect ACE2 expression in the lung and associated respiratory tissue.

Age and male sex are significant risk factors for severe SARS-CoV-2 infections^13,29^. We therefore investigated whether either feature was associated with increased ACE2 expression. In four different cohorts of aging mice and one cohort of aging rats, we did not detect a significant age-dependent change in ACE2 expression in the lung (Figure 1A-B and S1D-F)^25,30–33^. Similarly, ACE2 expression in rodent lungs was not significantly different between sexes (Figure 1C-D)^25,34^. We next assessed the expression of ACE2 in three different human cohorts: 1) lung tissue from the Genotype-Tissue Expression project (GTEx)^23,24^, 2) whole-lung tissue samples from organ donors^35^, and 3) pathologically-normal lung tissue from a cohort of patients analyzed as part of The Cancer Genome Atlas^36^. These datasets yielded results that were consistent with our rodent analyses: ACE2 expression was equivalent between men and women and between young individuals (<29 years) and elderly individuals (>70 years)(Figure 1E-J). In total, these findings suggest that the increased morbidity of men and older patients with COVID-19 is unlikely to result from inherent differences in the basal level of ACE2 expression in the lung.

**Figure 1.**
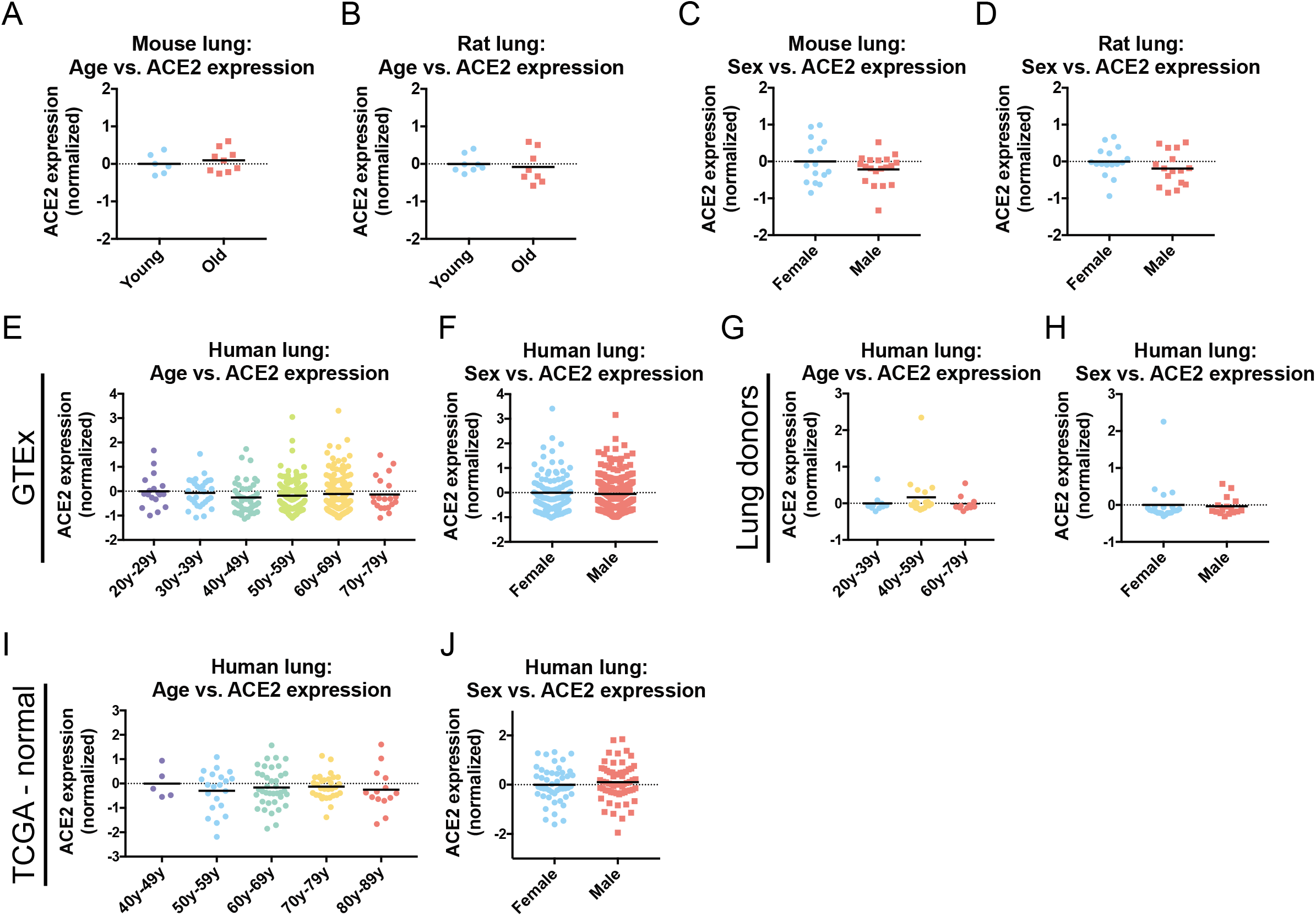
ACE2 expression in the lung is uncorrelated with age or sex. (A) ACE2 expression in the lungs of young mice (<26 weeks old) and old mice (>78 weeks old). (B) ACE2 expression in the lungs of young rats (6 weeks old) and old rats (104 weeks old). (C) ACE2 expression in the lungs of female mice and male mice. (D) ACE2 expression in the lungs of female rats and male rats. (E) ACE2 expression in lungs from the GTEx cohort by age. (F) ACE2 expression in lungs from the GTEx cohort by sex. (G) ACE2 expression in lungs from a cohort of organ donors by age. (H) ACE2 expression in lungs from a cohort of organ donors by sex. (I) ACE2 expression in pathologically normal lung tissue from patients from TCGA by age. (J) ACE2 expression in pathologically-normal lung tissue from patients from TCGA by sex. Each panel displays log2-normalized ACE2 expression relative to a control group. Data analyzed in A and C were from GSE132040. Data analyzed in B and D were from GSE53960. Data analyzed in E and F were from www.gtexportal.org. Data analyzed in G and H were from GSE1643. Data analyzed in I and J were from https://gdac.broadinstitute.org/. Additional information on the data sources and sample sizes are included in Table S1.

### Cigarette smoke increases the expression of ACE2 in the mammalian respiratory tract

Cigarette smoking is strongly associated with adverse outcomes from COVID-19^13,16,17,37–40^. To investigate whether smoking could affect ACE2 levels, we first assessed gene expression in mouse lungs. We analyzed a cohort of mice exposed to diluted cigarette smoke for 2, 3, or 4 hours per day for five months^41^. Strikingly, we found a dose-dependent increase in ACE2 expression according to smoke exposure (Figure 2A). Mice exposed to the highest dose of cigarette smoke expressed ~80% more ACE2 in their lungs compared to sham-treated mice. To determine whether this association was present in humans as well, we assessed tissue collected from three cohorts of current smokers and never-smokers^42–44^. For these analyses, lung epithelial cells were sampled by fiberoptic bronchoscopy from either the trachea, the large airways, or the small airways (Figure 2B). In each cohort, we observed that tissue samples from smokers exhibited ~30%-55% more ACE2 compared to tissue from non-smokers (Figure 2C-E). In a combined analysis comprised of data from all three tissue locations, ACE2 was in the top 2% of genes most strongly dysregulated by smoke exposure (Figure 2F).

**Figure 2.**
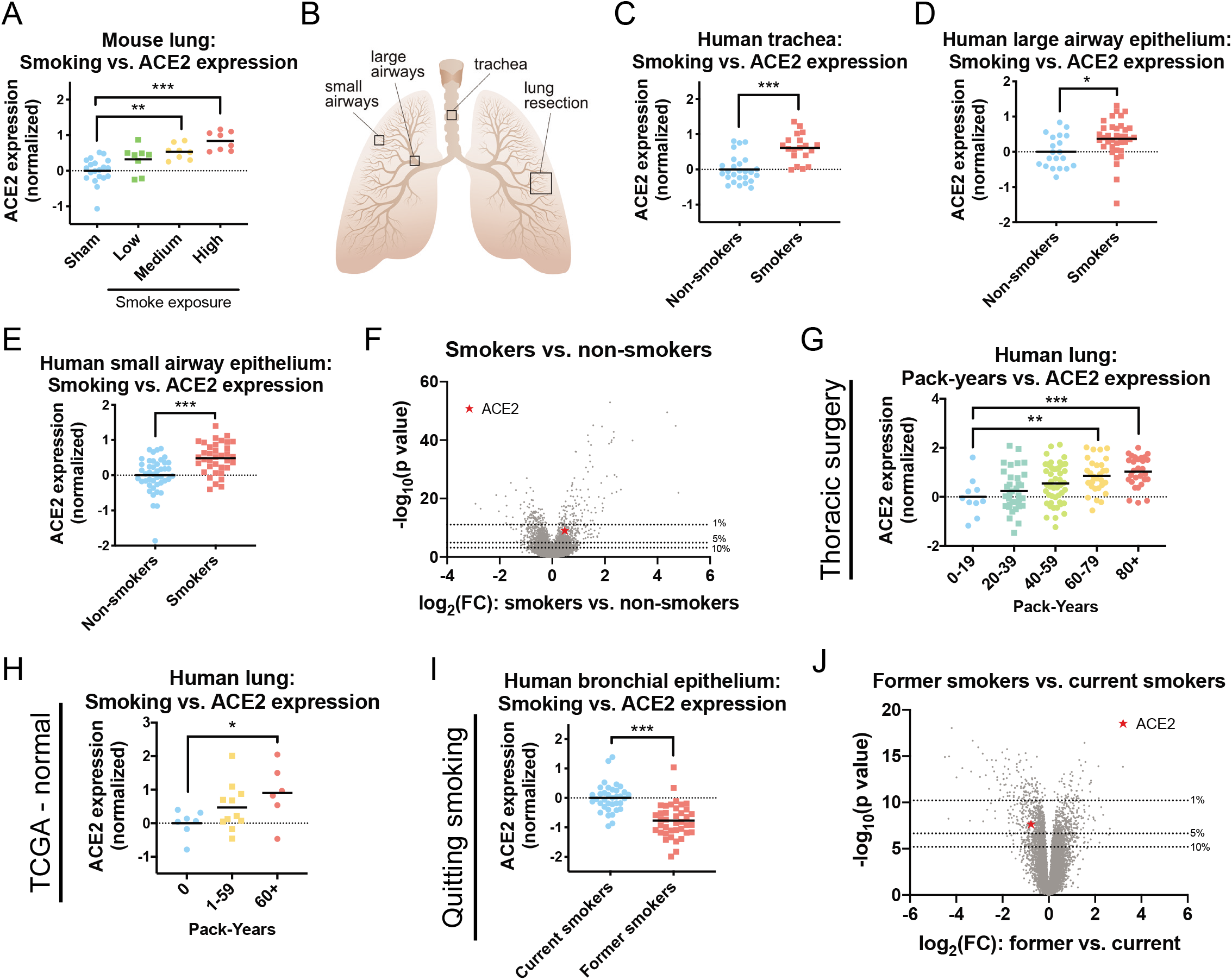
Cigarette smoke increases the expression of ACE2 in mouse and human lungs. (A) ACE2 expression in the lungs of mice that were sham-treated or that were exposed to diluted cigarette smoke for two, three, or four hours a day (low, medium, and high smoke exposure, respectively). (B) A diagram showing the approximate locations of tissue samples used in this analysis. Tracheal, large airway epithelial, and small airway epithelial specimens were collected by fiberoptic bronchoscopy as described in their respective publications. Lung resections were collected surgically from various locations. (C) ACE2 expression in human tracheal epithelia analyzed according to smoking history. (D) ACE2 expression in human large airway epithelia analyzed according to smoking history. (E) ACE2 expression in human small airway epithelia analyzed according to smoking history. (F) A volcano plot comparing gene expression in the respiratory epithelia of current smokers and never-smokers from C, D, and E. The dotted lines indicate various p-value thresholds (e.g., genes located above the 10^th^ percentile have a combined p value greater than 90% of the genes included in the meta-analysis). The location of ACE2 is indicated with a red star. (G) ACE2 expression in the lungs of a cohort of patients undergoing thoracic surgery analyzed according to the number of pack-years each patient smoked. (H) ACE2 expression in the lungs of TCGA patients analyzed according to the number of pack-years each patient smoked. (I) ACE2 expression in respiratory epithelia collected by fiberoptic bronchoscopy among either current smokers or former smokers. (J) A volcano plot comparing gene expression between current smokers and former smokers. The dotted lines indicate various p-value thresholds (e.g., genes located above the 10^th^ percentile have a combined p value greater than 90% of the genes included in the metaanalysis). The location of ACE2 is indicated with a red star. Each panel displays log2-normalized ACE2 expression relative to a control group. Data analyzed in A were from GSE18344. Data analyzed in C were from GSE13933. Data analyzed in D were from GSE22047. Data analyzed in E were from GSE64614. Data analyzed in G were from GSE76925. Data analyzed in H were from https://gdac.broadinstitute.org/. Data analyzed in I were from GSE79209. Additional information on the data sources and sample sizes are included in Table S1. *, p < .05; **, p < .005; ***, p < .0005 (Student’s t-test).

Next, we sought to determine whether human ACE2 expression showed a dose-dependent relationship with cigarette smoke, as we had observed in mice. To investigate this, we analyzed two human datasets: 1) lung tissue from a cohort of smokers undergoing thoracic surgery for transplantation, lung volume reduction, or nodule resection^45^ and 2) pathologically-normal lung tissue from TCGA patients^36^. In both cohorts, lung samples from patients who reported smoking the greatest number of pack-years also expressed the highest levels of ACE2 (Figure 2G-H). For instance, among smokers undergoing thoracic surgery, patients who had smoked more than 80 pack-years exhibited a ~100% increase in ACE2 expression relative to patients who had smoked less than 20 pack-years (Figure 2I).

We then investigated whether other demographic features could explain the upregulation of ACE2 in the lungs of smokers. Multivariate linear regression on the thoracic surgery cohort confirmed that smoking history was a significant predictor of ACE2 expression even when controlling for a patient’s age, sex, race, and body-mass index (Table 1). In the TCGA cohort, pack-year history did not remain significantly associated with ACE2 in a regression with additional clinical variables, which may reflect the very small number of patients who could be included in this analysis (22 patients; data not shown). However, demographic information was available for the tracheal epithelium cohort analyzed above (Figure 2E), and smoking status remained correlated with ACE2 in a regression that included each patient’s age, sex, and race (Table 1).

**Table 1.** Patient characteristics and ACE2 expression. Multivariate regression between various patient characteristics, smoking status, and ACE2 expression. The thoracic surgery cohort is from GSE76925 and the tracheal epithelium cohort is from GSE13933. Bolding indicates significant variables. *, p < .05; **, p < .005; ***, p < .0005.

|  |  | **β value** | **Standard Error** | **P value** |
| --- | --- | --- | --- | --- |
| **Thoracic surgery cohort** | **Age** | 0.0014 | 0.0086 | 0.8700 |
|  | **BMI** | -0.0199 | 0.0127 | 0.1193 |
|  | **Smoking Pack-years** | 0.0081 | 0.0025 | **0.0015\*\*** |
|  | **Race**<br>Caucasian = 0<br>Not Caucasian = 1 | 0.1910 | 0.1663 | 0.2527 |
|  | **Sex**<br>Female = 0<br>Male = 1 | 0.3335 | 0.1338 | **0.0138\*** |
| **Tracheal epithelium cohort** | **Age** | -0.0027 | 0.0080 | 0.7364 |
|  | **Smoking status**<br>Never smoker = 0<br>Current smoker = 1 | 0.5638 | 0.1269 | **7 x 10<sup>-5</sup>\*\*\*** |
|  | **Race**<br>Caucasian = 0<br>Not Caucasian = 1 | 0.1252 | 0.1331 | 0.3527 |
|  | **Sex**<br>Female = 0<br>Male = 1 | 0.2538 | 0.1328 | 0.0632 |

Finally, we examined the effects of quitting smoking on ACE2 expression. In a cohort of patients comprised of either current smokers or former smokers who had refrained from smoking for at least 12 months, quitting smoking was associated with a ~40% decrease in ACE2 expression (Figure 2I). ACE2 was among the top 5% of genes most strongly affected by quitting smoking (Figure 2J). In total, our results demonstrate that exposure to cigarette smoke increases the expression of the coronavirus receptor ACE2 in rodent and human respiratory tissue, and this upregulation is potentially reversible.

Coronavirus infections are facilitated by a set of host proteases that cleave and activate the viral spike (S) protein^46^. SARS-CoV-2 primarily relies on the serine protease TMPRSS2 but can also utilize an alternate pathway involving Cathepsin B/L in TMPRSS2-negative cells^6^. Interestingly, we observed that Cathepsin B expression, but not TMPRSS2 or Cathepsin L expression, was consistently increased in mice and humans exposed to cigarette smoke (Figure S3). In a metaanalysis across the trachea, large airways, and small airways, Cathepsin B was in the top 11% of genes dysregulated in the respiratory tract of cigarette smokers (Figure S3D). Thus, smoking can upregulate both the coronavirus receptor as well as a protease that SARS-CoV-2 is capable of using for viral activation.

### ACE2 is expressed at high levels in secretory cells in the lung epithelium

Mammalian lungs harbor more than 30 distinct cell types representing a variety of epithelial, endothelial, stromal, and immune compartments^47^. Of note, the upper respiratory epithelium is comprised of mucociliary cells, including goblet cells, club cells, and ciliated cells, that secrete protective fluids and remove inhaled particles from the airways^48^. The lower respiratory epithelium includes alveolar type 1 cells, which allow gas exchange with the blood, and alveolar type 2 cells, which regulate alveolar fluid balance and can differentiate into type 1 cells following injury^49^.

To gain further insight into coronavirus infections, we profiled multiple single-cell RNA-Seq experiments to identify the cell type(s) that express ACE2. We first examined a dataset containing 13,822 cells from normal mouse lungs^50^. We performed unsupervised Leiden clustering to separate the cells into distinct populations and then we assigned cell types to major clusters using established markers^51–54^. ACE2 was expressed solely in the EpCAM+ clusters that comprise the lung epithelium^55^, and was not detected in CD45+ immune cells^56^, PDGFRA+ mesenchymal cells^57^, or TMEM100+ endothelial cells (Figure 3A-B)^58^. We therefore focused on localizing ACE2 within the epithelial lineage. We found that ACE2 was present in a cluster of cells that express secretory markers including MUC5AC, GABRP, and SCGB1A1 that we identified as being comprised of closely-related goblet and club cells (Figure 3C-D)^54,59–61^. ACE2 expression was also observed in a subset of LAMP3+ alveolar type 2 cells^62^, but was largely absent from RTKN2+ alveolar type 1 cells^54^.

**Figure 3.**
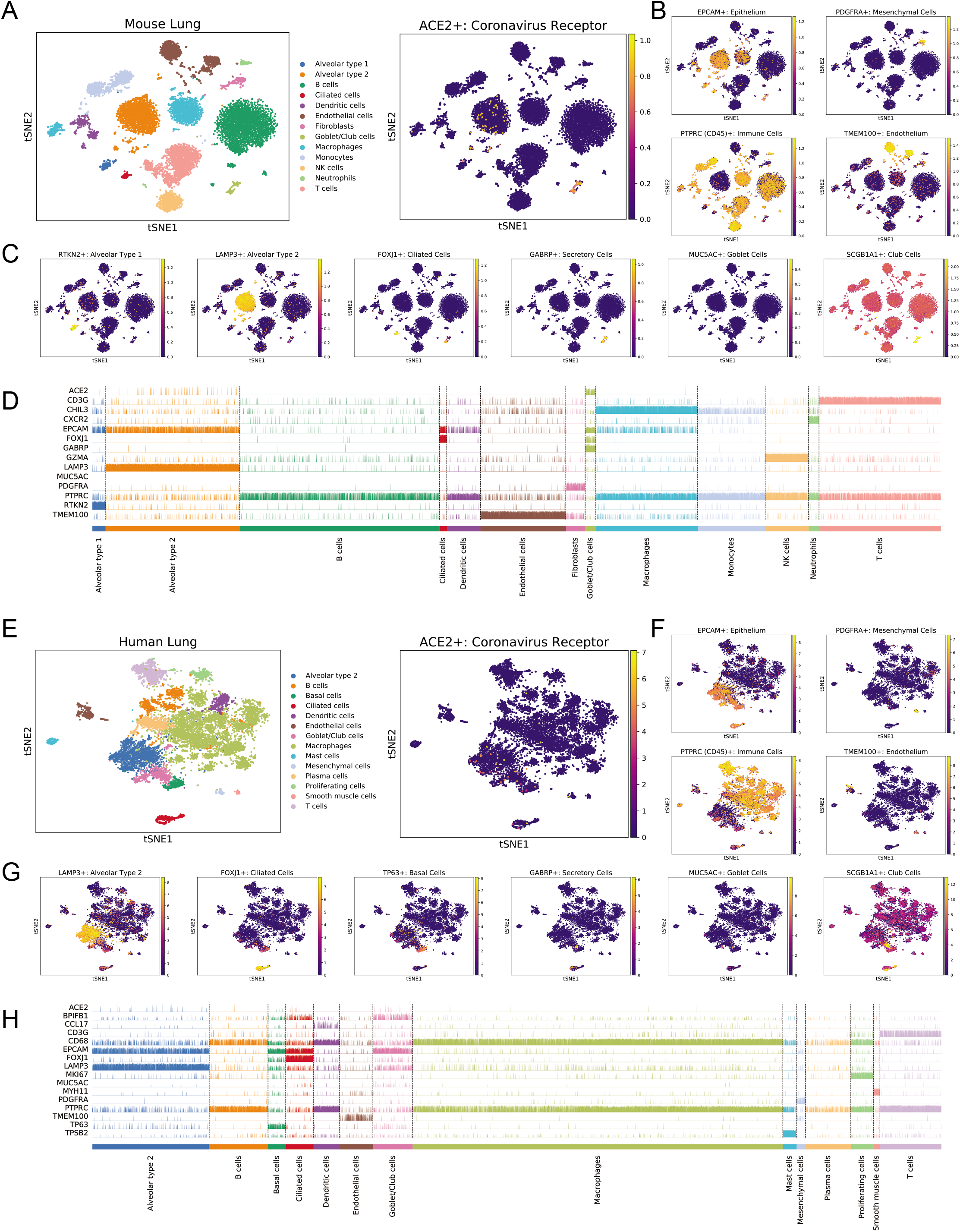
ACE2 is expressed in secretory club and goblet cells along with alveolar type 2 cells in the mammalian lung. (A) T-SNE clustering of cells from the mouse lung. Cells expressing ACE2 are highlighted in the right panel. (B) Cells in the mouse lung that express various lineage markers (TMEM100 for endothelial cells, EPCAM for epithelial cells, PDGFRA for mesenchymal cells, and PTPRC for immune cells) are highlighted. (C) Cells expressing markers for various epithelial lineages are highlighted: RTKN2 for alveolar type 1 cells, LAMP3 for alveolar type 2 cells, FOXJ1 for ciliated cells, GABRP for both goblet and club cells, MUC5AC for goblet cells, and SCGB1A for club cells. (D) A track plot displaying the expression of ACE2 and several lineage-related genes in different cell populations obtained from Leiden clustering. (E) T-SNE clustering of cells from the human lung. Cells expressing ACE2 are highlighted in the right panel. (F) Cells in the human lung that express various lineage markers (TMEM100 for endothelial cells, EPCAM for epithelial cells, PDGFRA for mesenchymal cells, and PTPRC for immune cells) are highlighted. (G) Cells expressing markers for various epithelial lineages are highlighted: LAMP3 for alveolar type 2 cells, FOXJ1 for ciliated cells, TP63 for basal cells, GABRP for both goblet and club cells, MUC5AC for goblet cells, and SCGB1A1 for club cells. (H) A track plot displaying the expression of ACE2 and several lineage-related genes in different cell populations obtained from Leiden clustering. The gene expression data used in A, B, C, and D are from GSE121611. The gene expression data used in D, E, F, and G are from GSE122960. Additional information on the data sources and sample sizes are included in Table S1.

We then extended our findings to an independent single-cell dataset from human lungs^50^. Consistent with our initial observations, we found that ACE2+ cells were found almost exclusively in the EPCAM+ epithelial compartment (Figure 3E-F). Within this lineage, we observed ACE2 expression in the LAMP3+ alveolar type 2 cluster and the MUC5AC+ goblet/club cell cluster (Figure 3G-H). Additionally, in human cells, there was some expression of ACE2 in FOXJ1+ ciliated cells^63^. In total, our combined analysis demonstrates that ACE2 is expressed in the mammalian lung epithelium, and is present at particularly high levels in secretory club and goblet cells as well as in alveolar type 2 cells.

### Cigarette smoke triggers an increase in ACE2+ cells by driving secretory cell expansion

Our findings on ACE2’s expression pattern suggested a possible explanation for its upregulation in the lungs of cigarette smokers. Chronic exposure to cigarette smoke has been reported to induce the expansion of secretory goblet cells, which produce mucous to protect the respiratory tract from inhaled irritants^64–66^. Thus, the increased expression of ACE2 in smokers’ lungs could be a byproduct of smoking-induced secretory cell hyperplasia.

To investigate this hypothesis, we examined a dataset of single-cell transcriptomes collected from the tracheas of current smokers and never-smokers^67^. As these cells were derived from the upper airways, they were highly-enriched for pseudostratified respiratory epithelial cell types and contained few alveolar cells. Consistent with the initial analysis of this dataset, Leiden clustering revealed a large population of TP63+ basal cells (the putative stem cell population for the airway epithelium)^68^, an intermediate KRT8^high^ population that showed evidence of ongoing differentiation^67^, and populations of mature FOXJ1+ ciliated cells and MUC5AC+ secretory cells (Figure 4A-B). We also identified markers for submucosal gland (SMG) clusters as well as rare epithelial cell types, including CFTR+ ionocytes^69^ and CHGA+ neuroendocrine cells^70^. Within this dataset, ACE2 was expressed at the highest levels in the MUC5AC+ secretory cell cluster, followed by SMG-secretory cells, KRT8^high^ intermediate cells, and FOXJ1+ ciliated cells.

**Figure 4.**
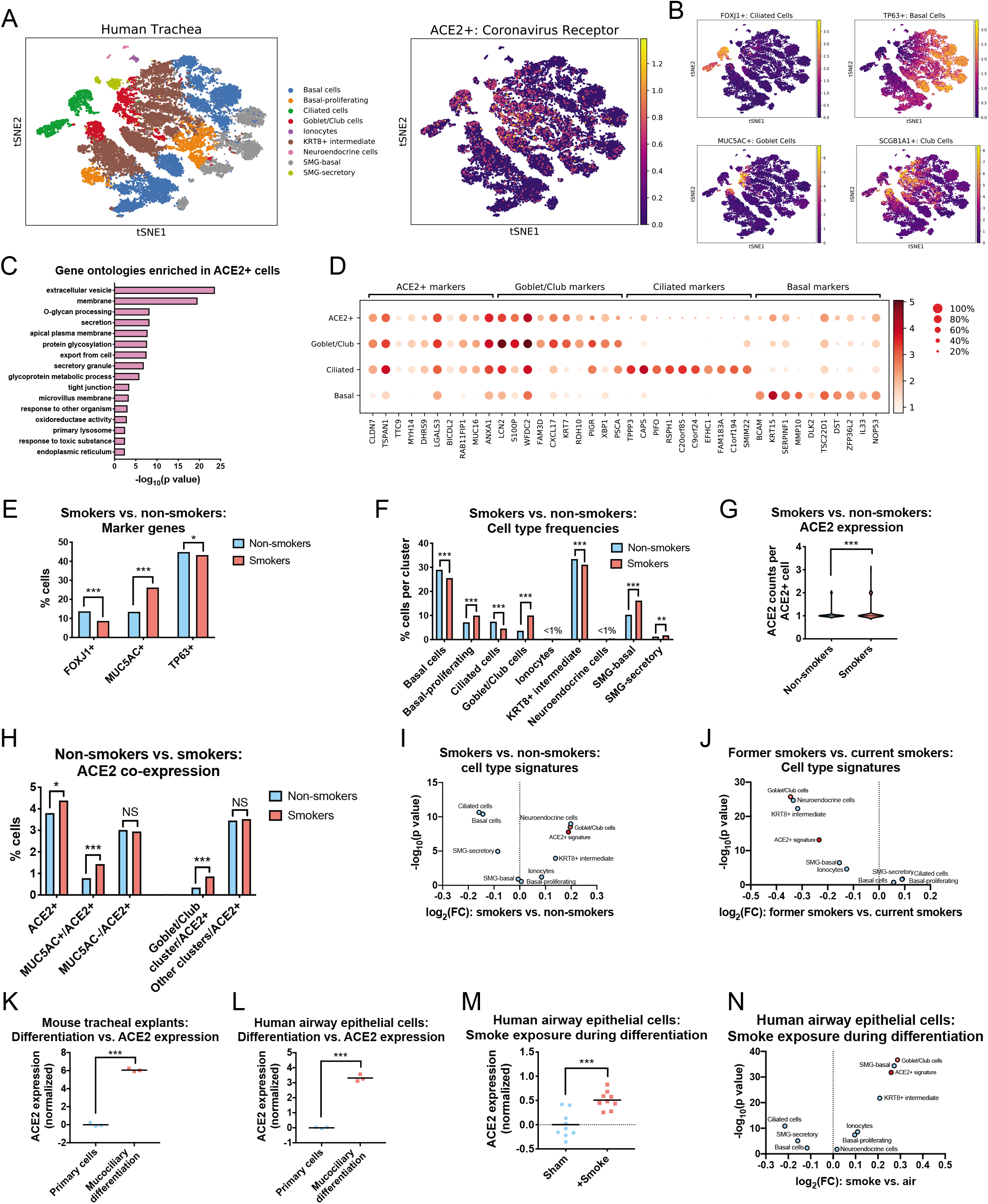
Cigarette smoke causes the expansion of ACE2+ secretory cells. (A) T-SNE clustering of the transcriptomes from single cells derived from the airway epithelia of smokers and never-smokers. Cells expressing ACE2 are highlighted in the right panel. (B) Cells in the human airway that express various lineage markers (FOXJ1 for ciliated cells, TP63 for basal cells, MUC5AC for goblet cells, and SCGB1A1 for club cells) are highlighted. (C) GO terms enriched among ACE2-correlated transcripts. (D) Dot plots displaying the expression of the top 10 differentially-expressed marker genes for various airway lineages and for ACE2+ cells. (E) The fraction of cells expressing the indicated marker genes are displayed. FOXJ1 is a marker for ciliated cells, MUC5AC is a marker for goblet cells, and TP63 is a marker for basal cells. (F) The fractions of cells found in each cell type cluster are displayed. (G) The numbers of counts per ACE2+ cell are displayed. (H) The fraction of ACE2+ cells co-expressing MUC5AC or that are found within the goblet/club cell cluster are displayed. (I) The 100 top-ranked differentially-expressed genes from each cluster in (A) as well as the 100 genes most strongly correlated with ACE2 were used to re-analyze the bulk gene expression data from smokers and non-smokers in Figure 2F. A volcano plot displays the mean expression change of each cell type signature. (J) The same transcriptional signatures as in Figure 4I were used to re-analyze the data from current smokers and former smokers in Figure 2J. A volcano plot displays the mean expression change of each cell type signature. (K) ACE2 expression in mouse tracheal explants undergoing mucociliary differentiation. (L) ACE2 expression in human airway epithelial cells undergoing mucociliary differentiation. (M) ACE2 expression in human airway epithelial cells that underwent mucociliary differentiation in the presence of clean air or cigarette smoke. (N) The same transcriptional signatures as in Figure 4I were used to re-analyze the data from smoke exposure during differentiation from Figure 4M. A volcano plot displaying the mean expression change of each cell type signature is displayed. Data analyzed in A through H were from GSE131391. Data analyzed in K were from GSE75715. Data analyzed in L were from GSE39059. Data analyzed in M were from GSE135188. Additional information on the data sources and sample sizes are included in Table S1. In E, F, and H, a chisquare test is applied. In G, a Mann-Whitney U test is applied. In K, L, and M, a Student’s t-test is applied. *, p < .05; **, p < .005; ***, p < .0005.

We next identified the transcripts whose expression correlated with ACE2. Across all cells, ACE2 levels were strongly correlated with several mucin genes, including MUC1, MUC4, MUC15, and MUC16, as well as other genes associated with barrier epithelia, including ALCAM^71^, CLDN7^72^, and TJP3^73^ (Table S2A). Gene ontology analysis revealed that ACE2-correlated transcripts were enriched for genes involved in secretion, glycosylation, and the response to toxic substances, consistent with the airway epithelium’s role as a producer of mucus and a barrier against foreign matter (Figure 4C and Table S2B)^74^. ACE2+ correlates overlapped with but were not identical to goblet/club markers: several genes were widely expressed in both populations (MUC16, CLDN7, S100P), while others were more restricted to single lineages (PIGR, PSCA)(Figure 4D). In general, the ACE2+ signature was expressed in both secretory cells and ciliated cells but not in the basal stem cell compartment.

We next separated the cells harvested from current smokers and never-smokers and then analyzed each population separately. Consistent with previous reports^64–66^, we detected a significant expansion of the secretory cell compartment in smokers’ airways, as evidenced by both cluster analysis and by the increase in the number of cells expressing the canonical goblet cell marker MUC5AC (Figure 4E-F). FOXJ1+ ciliated cells decreased in abundance, likely reflecting the fact that smoke exposure inhibits ciliogenesis^75^. Smoking caused an increase in both the number of ACE2+ positive cells and in ACE2 expression within ACE2+ cells (Figure 4G-H). This expansion resulted from hyperplasia of the secretory cell compartment: smoke exposure resulted in a two-fold increase in the frequency of MUC5AC+/ACE2+ double-positive cells, while there was no significant change in the frequency of MUC5AC-/ACE2+ cells. In total, these results provide single-cell evidence that the increase in ACE2 expression in smokers’ respiratory tracts is caused by the expansion of mucous-secreting goblet cells that co-express ACE2.

To further validate these findings, we derived gene signatures for each of the cell types identified from tracheal single-cell sequencing, and we applied these signatures to analyze the bulk gene expression datasets from smokers and non-smokers that we had previously investigated (Figure 2). The goblet/club cell gene signature was significantly upregulated in current smokers compared to never smokers, while in former smokers these genes were strongly downregulated (Figure 4I-J). A signature based on ACE2-correlated transcripts showed a similar expression pattern. In contrast, a ciliated cell transcriptional signature was strongly downregulated among current smokers. As some FOXJ1+ ciliated cells co-express ACE2, this smoking-dependent suppression of ciliogenesis may partially blunt the increase in ACE2+ cells caused by smoke-induced goblet cell hyperplasia.

Secretory cell differentiation of lung epithelium can be modeled *in vitro* by culturing cells at an air-liquid interface (ALI)^76,77^. Under appropriate conditions, primary respiratory cells growing at an ALI will undergo mucociliary differentiation into a stratified epithelium consisting of ciliated cells, goblet cells, and club cells^78^. As our single-cell analysis suggested that the coronavirus receptor ACE2 is expressed at higher levels in differentiated secretory and ciliated cells compared to basal stem cells, we investigated whether *in vitro* mucociliary differentiation increases ACE2 expression. Indeed, in mouse tracheal extracts^79^ and primary human lung cells^80^, mucociliary differentiation resulted in a highly-significant upregulation of ACE2 (Figure 4K-L). Finally, to investigate the link between smoking, differentiation, and ACE2 expression, we examined data from human bronchial epithelial cells cultured at an ALI in which cells were either exposed to clean air or to diluted cigarette smoke^81^. Remarkably, treatment with cigarette smoke during *in vitro* differentiation resulted in a significant upregulation of ACE2 relative to cells that were differentiated in clean air (Figure 4M). Smoke exposure increased ACE2 expression by ~42%, comparable to the increases that we observed between the lungs of non-smokers and smokers (Figure 2). Differentiation in the presence of cigarette smoke similarly resulted in an upregulation of the goblet/club cell transcriptional signature and a downregulation of the ciliated cell transcriptional signature (Figure 4N). In full, our results demonstrate that a subset of lung secretory cells express the coronavirus receptor ACE2, and cigarette smoke promotes the expansion of this cell population.

### ACE2 is upregulated in smoking-associated diseases and by viral infections

To follow up on these observations, we investigated whether ACE2 expression was affected by other lung diseases and/or carcinogen exposures. Indeed, we observed increased ACE2 expression in multiple cohorts of patients with chronic obstructive pulmonary disease (COPD) and idiopathic pulmonary fibrosis (IPF)(Figure S3A-D)^82–85^. Interestingly, both COPD and IPF are strongly associated with prior cigarette exposure^86,87^, and COPD in particular has been identified as a risk factor for severe COVID-19^16,88^. However, ACE2 expression was generally not affected by other lung conditions or toxins. We did not observe a significant difference in ACE2 expression in lung samples from a large cohort of patients with asthma or from patients with the lung disease sarcoidosis (Figure S3E-F)^89,90^. Similarly, ACE2 expression was unaltered in lung tissue from a mouse model of cystic fibrosis and in mice exposed to a variety of carcinogens, including arsenic, ionizing radiation (IR), and 1,3-butadiene (Figure S3G-J)^91–94^. We conclude that ACE2 upregulation in the lung is tightly associated with a history of cigarette smoking and is not a universal response to pulmonary diseases.

So-called “cytokine storms”, characterized by high levels of circulating inflammatory cytokines, have been identified as a cause of COVID-19-related mortality^18,20,21^. Cytokine release can be triggered by viral infections, which serve to induce immune cell activation and expansion^95^. Cigarette smoke is also an inflammatory agent, and smokers tend to exhibit an increase in inflammation-related serological markers^96,97^. To investigate a potential link between inflammation and the expression of the host factors required for coronavirus infections, we first examined the levels of ACE2 in published datasets of respiratory epithelial cells challenged with different viruses. We observed a highly significant upregulation of ACE2 in airway cells that were infected with two RNA viruses, influenza and Respiratory Syncytial Virus (Figure 5A-B)^98^. Infection with two coronaviruses, SARS and MERS, that are closely related to the 2019 pandemic strain also caused an increase in ACE2 levels (Figure 5C-D)^99,100^. Double-stranded RNA that is produced during viral replication can be detected by host pattern recognition receptors, triggering a strong immune response^101^. Accordingly, transfection of the dsRNA mimic poly(I:C) into airway epithelial cells was sufficient to induce the upregulation of ACE2 (Figure 5E)^102^. These results reveal that inflammatory signals – like those triggered by smoking or by a viral infection – are capable of increasing the expression of ACE2 in the respiratory epithelium.

**Figure 5.**
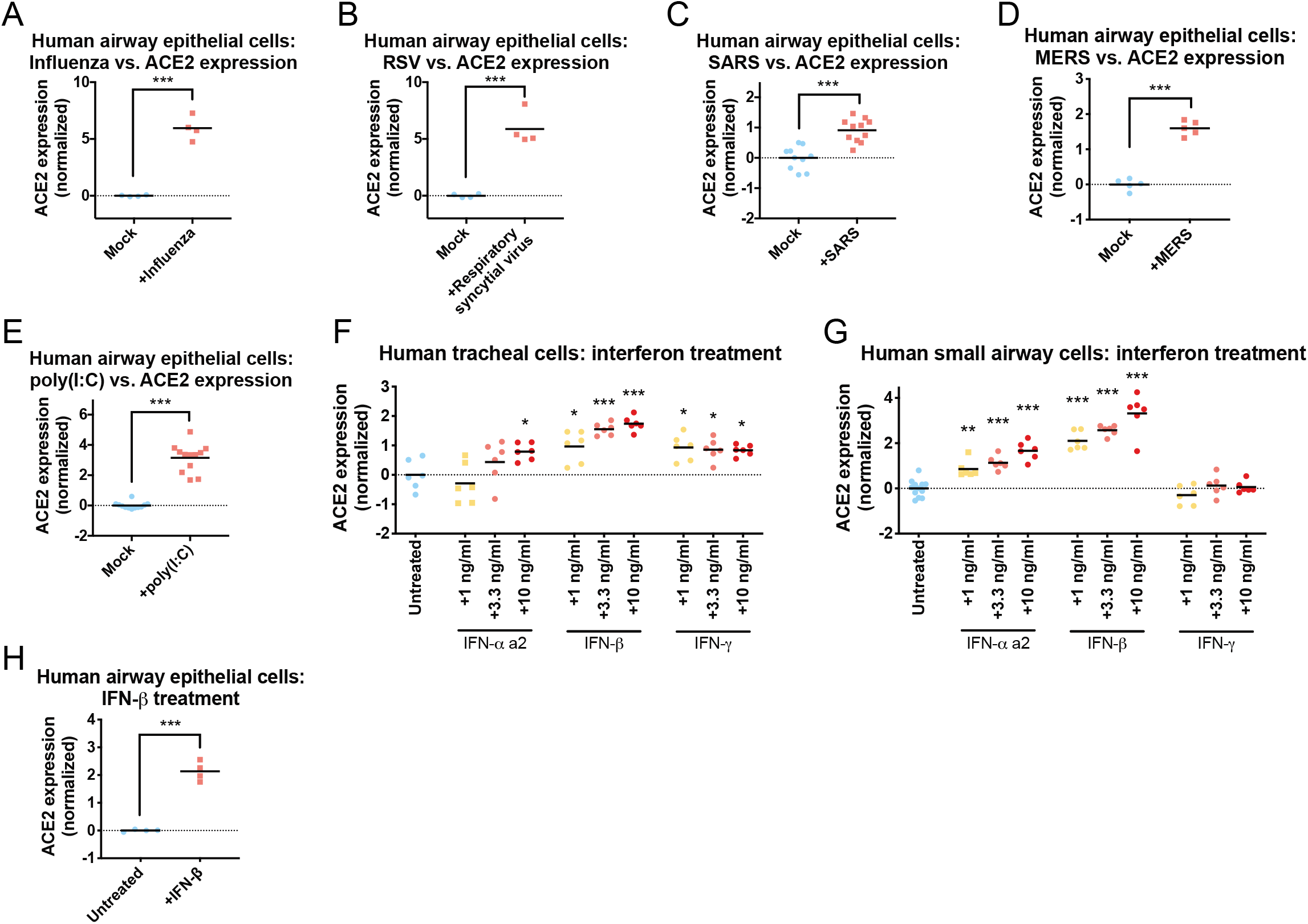
ACE2 is an interferon-stimulated gene that is upregulated by viral infections. (A) ACE2 expression in airway epithelial cells that were infected with influenza. (B) ACE2 expression in airway epithelial cells that were infected with respiratory syncytial virus. (C) ACE2 expression in airway epithelial cells that were infected with SARS. (D) ACE2 expression in airway epithelial cells that were infected with MERS. (E) ACE2 expression in airway epithelial cells that were transfected with the dsRNA mimic poly(I:C). (F) ACE2 expression in human tracheal cells that were cultured in the presence of the indicated cytokine for 24 hours. (G) ACE2 expression in human small airway epithelial cells that were cultured in the presence of the indicated cytokine for 24 hours. (H) ACE2 expression in airway epithelial cells that were cultured in the presence of IFN-β. Each panel displays log2-normalized ACE2 expression relative to a control group. Data analyzed in A and B were from GSE32138. Data analyzed in C were from GSE47963. Data analyzed in D were from GSE100504. Data analyzed in E were from GSE51392. Data analyzed in H were from GSE19392. Additional information on the data sources and sample sizes are included in Table S1. *, p < .05; **, p < .005; ***, p < .0005 (Student’s t-test).

To further investigate the stimuli that are sufficient to upregulate ACE2, we cultured primary epithelial cells from either the small airways or the trachea in the presence of different compounds. We found that exposure to interferons resulted in a significant upregulation of ACE2 expression (Figure 5F-G). IFN-α, IFN-β, and IFN-γ treatment increased ACE2 expression in tracheal cells, while only IFN-α and IFN-β affected ACE2 in small airway cells. To further verify these results, we re-analyzed a published gene expression dataset of airway epithelial cells exposed to IFN-β, and we found a similar increase in ACE2 levels following interferon treatment (Figure 5H)^103,104^. The results identify ACE2 as an interferon-regulated gene and suggest a potential mechanism by which inflammatory stimuli could facilitate SARS-CoV-2 infections.

### ACE2 RNA and protein levels are tightly correlated

This paper describes a series of analyses and experiments to identify factors that drive the expression of the coronavirus receptor ACE2. However, these results could potentially be confounded if changes in ACE2 RNA levels fail to affect the steady-state levels of ACE2 protein. While we lack the data to directly test that link in the lungs of smokers, we conducted three additional analyses to investigate whether ACE2 RNA and protein expression are generally correlated. First, we examined a dataset of human cancer cell lines that had been profiled at the transcriptional level by RNA-Seq and at the protein level by mass spectrometry^105–107^. Across 53 cell lines, ACE2 RNA and protein levels were strongly correlated (r = 0.82, p < .0001; Figure S4A). Indeed, ACE2’s RNA-protein correlation coefficient was higher than 95% of human genes that could be reliably detected (Figure S4B). Next, we compared ACE2 RNA levels with ACE2 immunohistochemistry staining in untransformed human tissues. We similarly observed a strong correlation between ACE2 RNA and protein staining: five of the six tissues with the highest levels of RNA expression exhibited strong staining, while none of the 35 other tissues with lower RNA levels exhibited strong staining (Figure S4C)^108^. Lastly, we directly compared ACE2 expression in seven human cell lines by qRT-PCR and by western blotting. We found that cell lines that displayed the highest levels of ACE2 RNA (LoVo, Caco-2, and Calu3) also exhibited the most ACE2 protein, while cell lines with very low levels of ACE2 RNA (HCT116, CAL148, and A2780) had undetectable levels of ACE2 protein (Figure S4D-E). Thus, while we cannot rule out the possibility that ACE2 RNA and protein levels are differentially regulated in certain circumstances, our analysis reveals that ACE2 RNA and protein levels are generally correlated.

## Discussion

While SARS-CoV-2 has infected more than 3 million people worldwide, fewer than 5% of COVID-19 cases are fatal^4,13^. Here, we show that cigarette smokers harbor consistently higher levels of the SARS-CoV-2 receptor ACE2 in their respiratory tracts. This upregulation is likely mediated by the expansion of ACE2+ secretory cells caused by chronic smoke exposure. Certain inflammatory cytokines also trigger ACE2 upregulation, which could further influence ACE2 expression due to smoking-associated lung inflammation. The overabundance of ACE2 in the lungs of smokers may partially explain why smokers are significantly more likely to develop severe SARS-CoV-2 infections that require aggressive medical interventions^13,16,17,37^. Furthermore, as quitting smoking is associated with a decrease in ACE2 expression, it is possible that giving up cigarettes may reduce susceptibility to deadly COVID-19.

Several contrasting findings exist in the literature on ACE2 and cigarette exposure^109–116^. In particular, it has been reported that nicotine and/or cigarette smoke has the potential to downregulate ACE2 expression in certain tissues or cell types^109–112^. In this manuscript, we focused our analysis on factors affecting ACE2 expression in the mammalian lungs and associated respiratory epithelia. We observed a consistent correlation between smoking history and ACE2 expression that was dose-dependent (Figure 2G-H), that could be recapitulated in mice (Figure 2A) and *in vitro* (Figure 4M), that was detectable in both bulk and single-cell analyses (Figure 2F and 4G-H), and that remained significant when controlling for other demographic variables (Table 1). Thus, we propose that cigarette smoke causes the upregulation of ACE2 expression in the respiratory tract. However, we recognize that individual components found within cigarettes, like nicotine, may have a different effect on ACE2 than whole smoke, and smoking itself could alter ACE2 levels in non-respiratory organs in different ways^110^.

Several previous studies have examined the localization of ACE2 within the respiratory tract^116–127^. Consistent with our results, ACE2 expression has been detected in the respiratory epithelium, including prominent staining in alveolar type 2 cells. However, some prior studies have also documented ACE2 expression in lung endothelial and smooth muscle cells^118,120,121^, which we did not observe. These findings could represent non-specific staining resulting from the particular antibodies that were used^128^. Alternately, ACE2 may be expressed in mesenchymal or other lineages within the lung, and our inability to detect it may be a limitation of the single-cell samples that we analyzed. Nonetheless, by examining datasets from both mice and humans, and by including cells collected from bronchial brushing, we demonstrated consistently high levels of ACE2 transcripts in the secretory cells that participate in mucociliary clearance^48^. As these cells line the upper respiratory tract, they may represent the initial site of coronavirus infections, followed by an eventual spread and migration into the alveoli.

Cigarette smoke has a profound impact on the lungs of chronic smokers. Notably, prolonged smoke exposure triggers secretory cell hyperplasia, thereby increasing the production of mucous in the respiratory tract^64–66^. We found that this smoking-induced expansion of secretory cells also causes an increase in ACE2 levels, as smoking doubled the number of cells co-expressing ACE2 and MUC5AC. Additionally, consistent with a recent report^122^, we found that ACE2 is an interferon-regulated gene, and is over-expressed in lung epithelial cells following viral infection or interferon treatment. Lung damage and inflammation caused by smoking could also contribute to ACE2 upregulation. Additionally, we speculate that the interferon-dependent upregulation of ACE2 could create a positive-feedback loop for SARS-CoV-2 infections. That is, interferon secretion following an initial infection could increase ACE2 expression within neighboring cells, thereby rendering those cells susceptible to SARS-CoV-2 and facilitating viral dissemination. Accordingly, clinical interventions to dampen the immune response could benefit patients in part by breaking this positive-feedback cycle^22,129^.

The factors that mediate overall susceptibility to SARS-CoV-2 infections are poorly understood. We speculate that the increased expression of ACE2 that we found in the lungs of smokers could partially contribute to the severe cases of COVID-19 that have been observed in this patient population. Chronic smokers may exhibit a number of co-morbidities, including emphysema, atherosclerosis, and immune dysregulation^130^, that are also likely to affect COVID-19 progression.

While the effects of smoking can last for years, smoking cessation causes an improvement in lung function and an overall decrease in disease burden^130^. Quitting smoking leads to a normalization of respiratory epithelial architecture^131^, a decrease in hyperplasia^132^, and a downregulation of ACE2 levels. Thus, for multiple reasons, smoking cessation could eventually lessen the risks associated with SARS-CoV-2 infections.

### Limitations

Our analysis in this manuscript has a number of important limitations. First, the relationship between age, sex, and ACE2 expression remains controversial. In the literature, ACE2 expression in the lung has been reported to both increase^133^ and decrease^120,134^ during aging. Across five rodent and three human datasets, we did not observe a significant correlation in either direction between ACE2 levels and age. It remains possible that such a difference could be found using tissue from older or younger individuals. Additionally, it is possible that ACE2 expression within single cell types correlates with age, but such differences are not prominent enough to detect using bulk microarray and RNA-Seq analysis.

In order for SARS-CoV-2 to infect a host cell, the virus must bind to ACE2 protein that is present on the cell surface. Here, we have described the expression of ACE2 transcripts in mammalian lungs, but our work does not guarantee that stimuli affecting the levels of ACE2 mRNA will have the same effect on ACE2 protein. In general, we have documented a strong correlation between ACE2 mRNA and protein levels (Figure S4), but it remains possible that ACE2 protein is differentially regulated in mammalian lungs, or that the localization of ACE2 protein to the cell surface is strictly controlled. Immunohistochemistry using tissue samples from smokers and virus-infected patients will be needed to confirm that these factors also increase the levels of ACE2 protein available to interact with SARS-CoV-2 particles.

Finally, the exact role of ACE2 as a mediator of disease severity remains to be determined. Mice that were engineered to express high levels of human ACE2 succumbed to infections with the SARS coronavirus more quickly than mice that expressed low levels of human ACE2, suggesting that increasing ACE2 enhances viral susceptibility^9^. At the same time, ACE2-knockout mice are vulnerable to a variety of lung injuries, and ACE2 expression has been reported to play a protective role in the respiratory tract^8,135,136^. As ACE2 expression is both necessary and sufficient for SARS-CoV-2 infections^6,137^, it seems highly likely that an expansion of ACE2+ cells in the lungs will facilitate viral dissemination, but it remains possible that the presence of ACE2 has some beneficial functions as well. Additional work will be required to determine the precise impact of ACE2 expression levels on the clinical course of COVID19.

## Supporting information

Table S1 - data sources

Table S2 - GO terms

## Author contributions

Conceptualization, J.C.S. and J.M.S.; Methodology, J.C.S. and J.M.S.; Software, J.C.S.; Formal analysis, J.C.S. and J.M.S.; Writing – Original Draft, J.M.S.; Writing – Review & Editing, J.C.S. and J.M.S; Supervision, J.M.S.; Investigation, J.C.S., E.L.S., V.G., M.L.Y., K.M.J., and J.M.S..

## STAR Methods

### RESOURCE AVAILABILITY

Further information and requests for resources should be directed to and will be fulfilled by the Lead Contact, Dr. Jason Sheltzer.

### MATERIALS AVAILABILITY

This study did not generate any new unique reagents.

### DATA AND CODE AVAILABILITY

All of the data used in this manuscript are publicly accessible and are described in Table S1. All code for performing these analyses is available at github.com/joan-smith/covid19.

### EXPERIMENTAL MODELS AND SUBJECT DETAILS

#### Cell lines

All cell lines used in this study were acquired from ATCC, except for CAL148 cells that were acquired from DSMZ and A2780 cells that were acquired from ECACC. Primary small airway epithelial cells and primary bronchial/tracheal epithelial cells were cultured in Airway Epithelial Cell Basal Medium (ATCC; Cat. No. PCS-300-030) supplemented with the Bronchial Epithelial Cell Growth Kit (ATCC; Cat. No. PCS-300-040^™^). HCT116, CAL148, and A375 cell lines were cultured in Dulbecco’s Modified Eagle Medium (DMEM) (Gibco; Cat. No. 11995073) supplemented with 10% fetal bovine serum (FBS)(Corning; Cat. No. 35-010-CV). LoVo and A2780 were cultured in RPMI 1640 (Lonza; Cat. No. 12-115F) supplemented with 10% FBS. Calu-3 was cultured in Eagle’s Minimum Essential Medium (EMEM) (ATCC; Cat. No. 30-2003^™^) supplemented with 10% FBS. Caco-2 was cultured in EMEM supplemented with 20% FBS. All cell lines were grown in a humidified environment at 37°C and 5% CO2.

### METHOD DETAILS

#### Sample Sizes and Statistical Methodology

For the qPCR assays in Figure 5F-G and Figure S4D, we performed three biological replicates using two ACE2 primer pairs. Gene expression values in Figure 1, Figure S1, Figure 2, Figure S2, Figure 4K-M, Figure S3, and Figure 5 were compared by a two-sided Student’s t-test. Gene expression values in Figure 4G were compared by a Mann-Whitney U test. The fractions of cells expressing a particular marker in Figure 4E-F and Figure 4H were compared by a chi-square test. No outliers were excluded from these analyses. Sample sizes of the gene expression datasets that were analyzed are listed in Table S1.

#### Overall analysis strategy

The analysis described in this paper was performed using Python, Excel, and Graphpad Prism. Gene expression data was acquired from the Gene Expression Omnibus (GEO)^139^, the GTEx portal^23^, the Broad Institute TCGA Firehose^140^, the Human Cell Atlas^141^, and the Single-Cell Expression Atlas^142^, as described below. For microarray datasets, probeset definitions were downloaded from GEO, and probes mapping to the same gene were collapsed by averaging. For each gene expression comparison, a control population was identified (e.g., young rats, sham-treated mice, non-smokers, etc.), and gene expression values were log2-transformed and normalized by subtraction so that the mean expression of a gene of interest in the control population was 0. Graphs of gene expression values were then generated using Graphpad Prism; all data points are displayed and no outliers were excluded from analysis.

#### Data sources

The data sources used in this paper are listed in Table S1. In general, pre-processed microarray and RNA-Seq datasets were downloaded. Additional notes on sample selection and processing are included in Table S1.

#### Multivariate regression and smoking history

Multivariate regression to investigate the relationship between ACE2 expression and smoking history was performed on the GSE76925 lung tissue dataset and the GSE13933 tracheal epithelium dataset. Regressions were performed in Python using ordinary least squares from the statsmodels package^143^. Results reported include the standard errors (‘bse’), betas, and p-values.

#### Single-cell analysis

Single-cell clustering and analysis on the datasets listed in Table S1 was performed in Python using the Scanpy and Multicore-TSNE packages^51,144^. To filter out low-quality cells, only cells in which 500 or more genes were detected were included in this analysis. Before clustering, transcript counts were counts-per-million normalized and log2 transformed. Highly variable genes were selected using the Seurat approach in Scanpy, and these highly variable genes were used to produce the principal component analysis. A t-SNE projection and unsupervised Leiden clustering were then performed on each dataset using nearest neighbors, as described in the associated code. Parameter searches were performed according to the method described in ref^145^.

In order to label each cluster, a gene ranking analysis was obtained using Scanpy. The 20 most highly-ranked genes from each cluster (as determined by a t-test with overestimated variance) were identified. These genes were then compared against gold-standard marker lists from multiple sources to produce the cluster labels^51–54,67^.

#### Gene ontology analysis

In order to identify the genes whose expression correlates with ACE2, pairwise Pearson correlations coefficients (PCCs) were calculated between ACE2 and every other expressed gene. Genes whose PCC were more than three standard deviations greater than the average gene’s PCC were classified as strongly correlated with ACE2. Gene ontology terms enriched in this group were then identified with GProfiler against the background list of non-strongly correlated genes using a Benjamini-Hochberg FDR of .05^146^.

#### Volcano plots

For the meta-analysis of gene expression changes in the small airways, large airways, and trachea, p values from each independent dataset were combined using Fisher’s method. To deconvolve the bulk gene expression datasets based on cell type signatures, the 100 highest ranking genes from each cell type cluster (identified using a t-test with overestimated variance, as described above) were identified and the 100 genes exhibiting the strongest correlation with ACE2 were identified. Volcano plots were generated using the mean expression of the 100 genes comprising each signature and the p-value of a t-test comparing the signature genes to all genes within a dataset.

#### Cytokine treatments

Early-passage human tracheal epithelial cells and small airways epithelial cells were acquired from ATCC and cultured according to the supplier’s recommended method. Cells were grown for 48 hours to approximately 70% confluence, treated with the indicated compound for 24 hours, and then total RNA was harvested.

#### RNA expression analysis

qRT-PCR was performed as described in ref^147^. In short, RNA was extracted using TRIzol (Life Technologies; Cat. No. 15596018) and then purified with a Qiagen RNeasy Mini Kit (Cat. No. 74106). RNA was converted to cDNA using SuperScript IV VILO Master Mix (ThermoFisher Scientific; Cat. No. 11756500). Quantitative PCR was performed using SYBR Premier Ex Taq (Takara; Cat No. RR420L) and quantified using the QuantStudio 6 Flex Real-Time PCR system (Applied Biosystems). The primers for ACE2 were as follows: (set 1) 5’-CGAAGCCGAAGACCTGTTCTA and 5’-GGGCAAGTGTGGACTGTTCC and (set 2) 5’-CAAGAGCAAACGGTTGAACAC and 5’-CCAGAGCCTCTCATTGTAGTCT. GAPDH was assessed as a control using primers 5’-TGCACCACCAACTGCTTAGC and 5’-GGCATGGACTGTGGTCATGAG.

#### Western blotting

Western blotting was performed as described in ref^147^. In short, whole cell lysates were harvested and resuspended in RIPA buffer [25 mM Tris, pH 7.4, 150 mM NaCl, 1% Triton X 100, 0.5% sodium deoxycholate, 0.1% sodium dodecyl sulfate, protease inhibitor cocktail (Sigma; Cat. No. 4693159001), and phosphatase inhibitor cocktail (Sigma, Cat. No. 4906845001)]. Protein lysates were loaded onto a 10% SDS-PAGE gel. The Trans-Blot Turbo Transfer System (Bio-Rad) and polyvinylidene difluoride membranes were used for protein transfer. Antibody blocking was done with 5% milk in TBST (19 mM Tris base, NaCl 137 mM, KCl 2.7 mM and 0.1% Tween-20) for 1 hour at room temperature. Primary antibodies used were ACE2 (Abcam; Cat. No. ab108252), used at a 1:1000 dilution in 5% milk, and GAPDH (Santa Cruz Biotechnology; Cat. No. sc-365062), used at a 1:1000 dilution in 5% milk. Blots were incubated overnight at 4°C. Membranes were then washed three times for 10 minutes each before they were incubated in secondary antibodies for an hour at room temperature. For the ACE2 blot, goat anti-rabbit IgG H&L (HRP) (Abcam; Cat. No. ab-97051) was used as a secondary antibody at a 1:20000 dilution in 5% milk. For the GAPDH blot, goat anti-mouse IgG (H + L)-HRP Conjugate (Bio-Rad; Cat. No. 1706516) was used as a secondary antibody at a 1:20000 dilution in 5% milk. Membranes were washed three more times and then developed using ProtoGlow ECl (National Diagnostics; Cat. No. CL-300).

## Supplemental Figures and Tables

**Figure S1.**
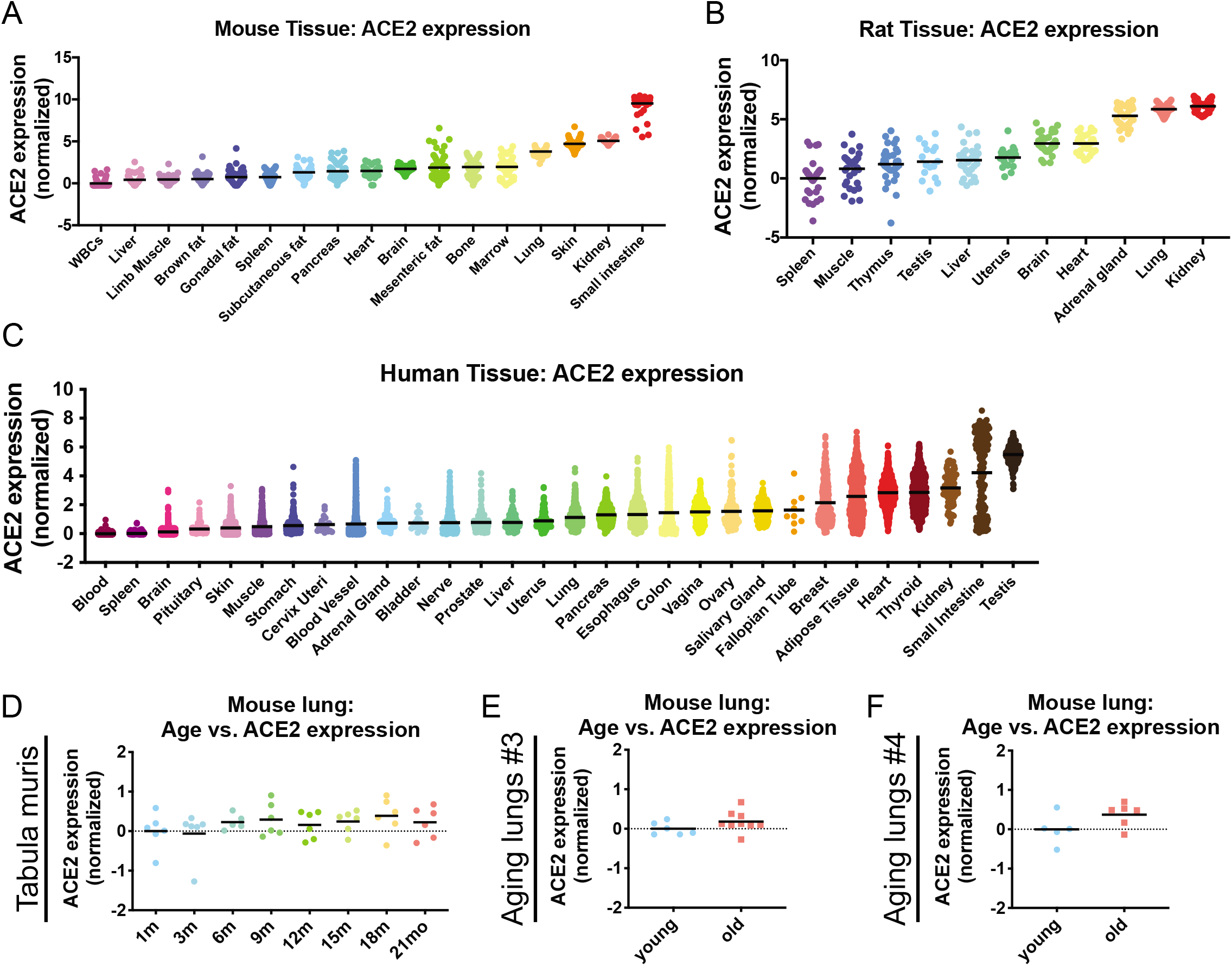
ACE2 expression varies between different mammalian tissues but is not strongly affected by aging. Related to Figure 1. (A) The expression of ACE2 in 17 different mouse tissues or cell types, sorted according to mean ACE2 expression. (B) The expression of ACE2 in 11 different rat tissues, sorted according to mean ACE2 expression. (C) The expression of ACE2 in 30 different human tissues or cell types, sorted according to mean ACE2 expression. (D) ACE2 expression in the lungs of a cohort of mice, sorted according to age. (E) ACE2 expression in the lungs of an independent cohort of young mice (2-month old) and old mice (18-month old and 26-month old). (F) ACE2 expression in the lungs of an independent cohort of young mice (7- to 9-week old) and old mice (14-month old). Each panel displays log2-normalized ACE2 expression relative to a control group. Data analyzed in A and D were from GSE132040. Data analyzed in B were from GSE53960. Data analyzed in C were from www.gtexportal.org. Data analyzed in E were from GSE6591. Data analyzed in F were from GSE80680. Additional information on the data sources and sample sizes are included in Table S1.

**Figure S2.**
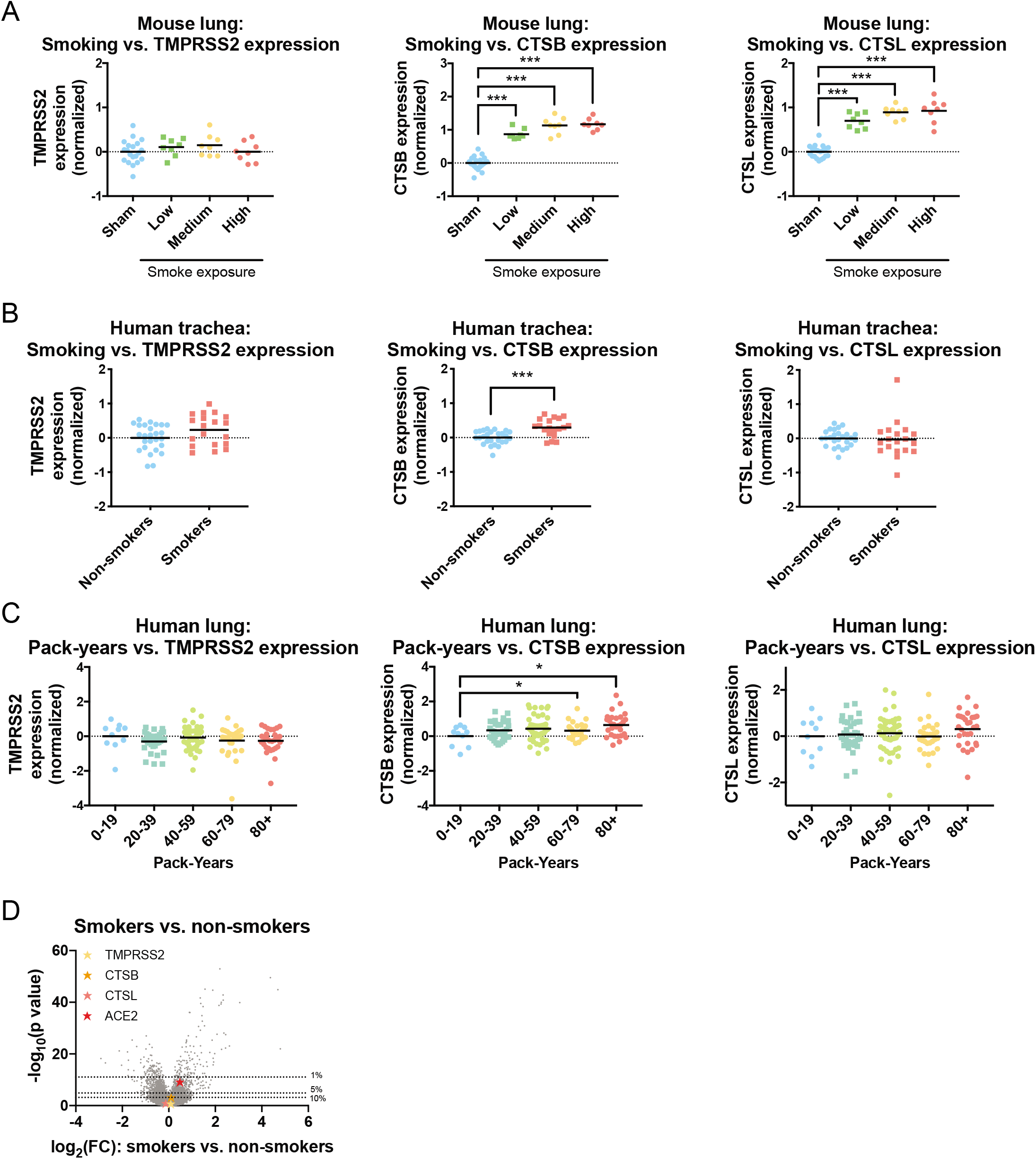
Cigarette smoke increases the expression of the coronavirus-associated protease Cathepsin B. Related to Figure 2. (A) The expression of TMPRSS2, Cathepsin B, and Cathepsin L in the lungs of mice that were sham-treated or that were exposed to diluted cigarette smoke for two, three, or four hours a day (low, medium, and high smoke exposure, respectively). (B) The expression of TMPRSS2, Cathepsin B, and Cathepsin L in human trachea collected by fiberoptic brushing and analyzed according to smoking history. (C) The expression of TMPRSS2, Cathepsin B, and Cathepsin L in human lung tissue in a cohort of smokers and analyzed according to the number of pack-years smoked. (D) A volcano plot comparing gene expression in the respiratory epithelia of current smokers and never-smokers from Figure 2C, 2D, and 2E. The dotted lines indicate various p-value thresholds (e.g., genes located above the 10^th^ percentile have a combined p value greater than 90% of the genes included in the meta-analysis). Each panel displays log2-normalized expression of the indicated gene relative to a control group. Data analyzed in A were from GSE18344. Data analyzed in B were from GSE13933. Data analyzed in C were from GSE76925. Additional information on the data sources and sample sizes are included in Table S1. *, p < .05; **, p < .005; ***, p < .0005 (Student’s t-test).

**Figure S3.**
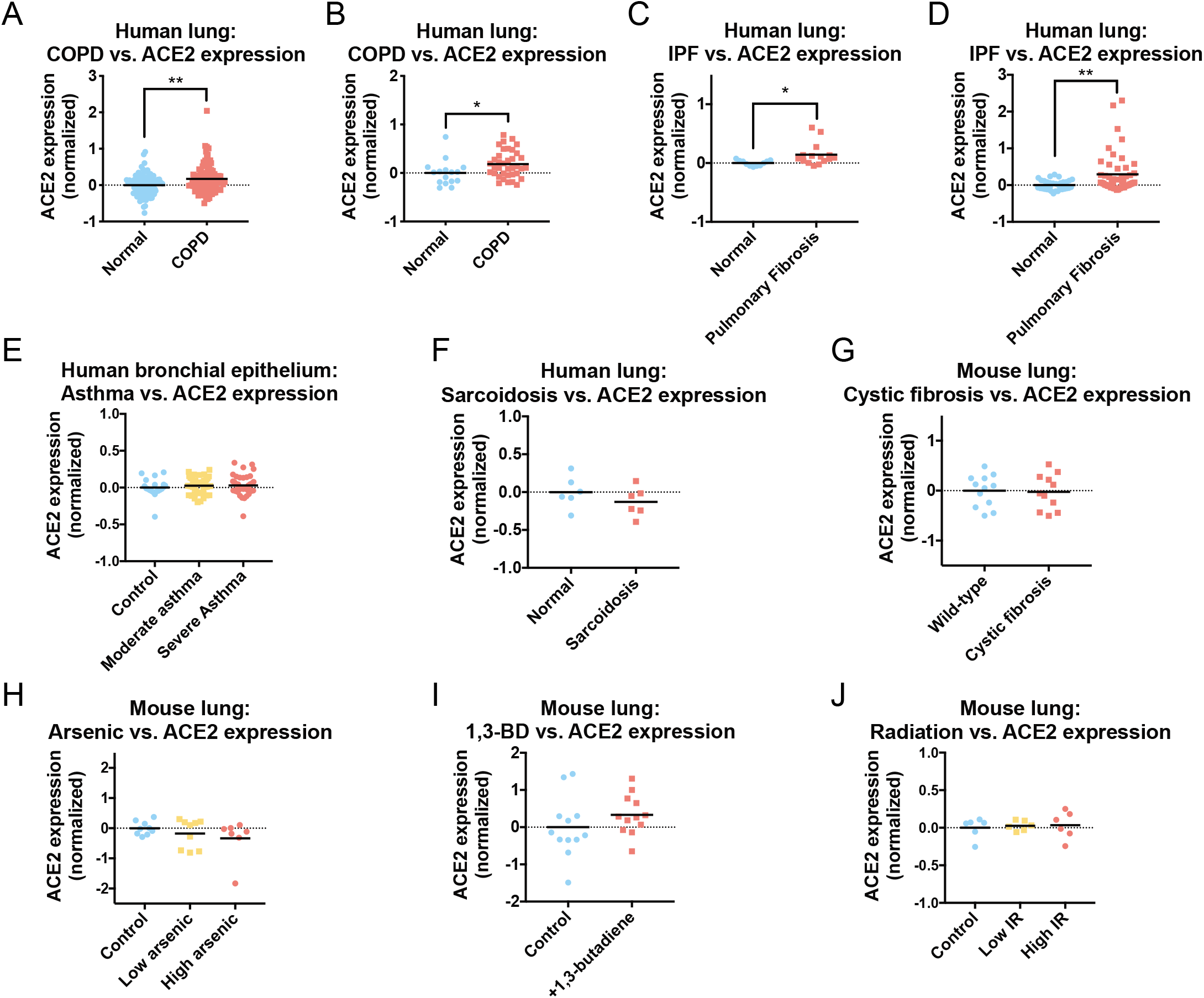
ACE2 is not commonly upregulated by lung diseases or carcinogen exposure. Related to Figure 4. (A) ACE2 expression in normal lung tissue and in lung tissue from patients with COPD. (B) ACE2 expression in normal lung tissue and in lung tissue from patients with COPD. (C) ACE2 expression in lung tissue from patients with idiopathic pulmonary fibrosis or from histologically-normal lung tissue from patients with lung cancer. (D) ACE2 expression in tissue from the lungs of transplant recipients with idiopathic pulmonary fibrosis or from control donor lungs. (E) The expression of ACE2 in bronchial brushing from individuals with normal lung function, mild asthma, or severe asthma. (F) The expression of ACE2 in disease-free lung tissue resected from patients with sarcoidosis or control tissue from healthy lung donors. (G) The expression of ACE2 in the lungs of a cystic fibrosis mouse model (*Cftr*^-/-^) or wild-type littermate controls. (H) The expression of ACE2 in the lungs of mice provided with normal water, water with 10 parts per billion sodium arsenite, or water with 100 parts per billion sodium arsenite (low and high arsenic, respectively). (I) The expression of ACE2 in the lungs of mice exposed to normal air or exposed to the carcinogen 1,3-butadiene. (J) The expression of ACE2 in the lungs of mice that were sham-treated, exposed to 5 Gy of ionizing radiation to the thorax, or exposed to 17.5 Gy of ionizing radiation to the thorax (low and high IR, respectively). Each panel displays log2-normalized ACE2 expression relative to a control group. Data analyzed in A were from GSE57148. Data analyzed in B were from GSE103174. Data analyzed in C were from GSE2052. Data analyzed in D were from GSE47460. Data analyzed in E were from GSE43696. Data analyzed in F were from GSE16538. Data analyzed in G were from GSE3100. Data analyzed in H were from GSE11056. Data analyzed in I were from GSE86623. Data analyzed in J were from GSE41789. Additional information on the data sources and sample sizes are included in Table S1. *, p < .05; **, p < .005; ***, p < .0005 (Student’s t-test).

**Figure S4.**
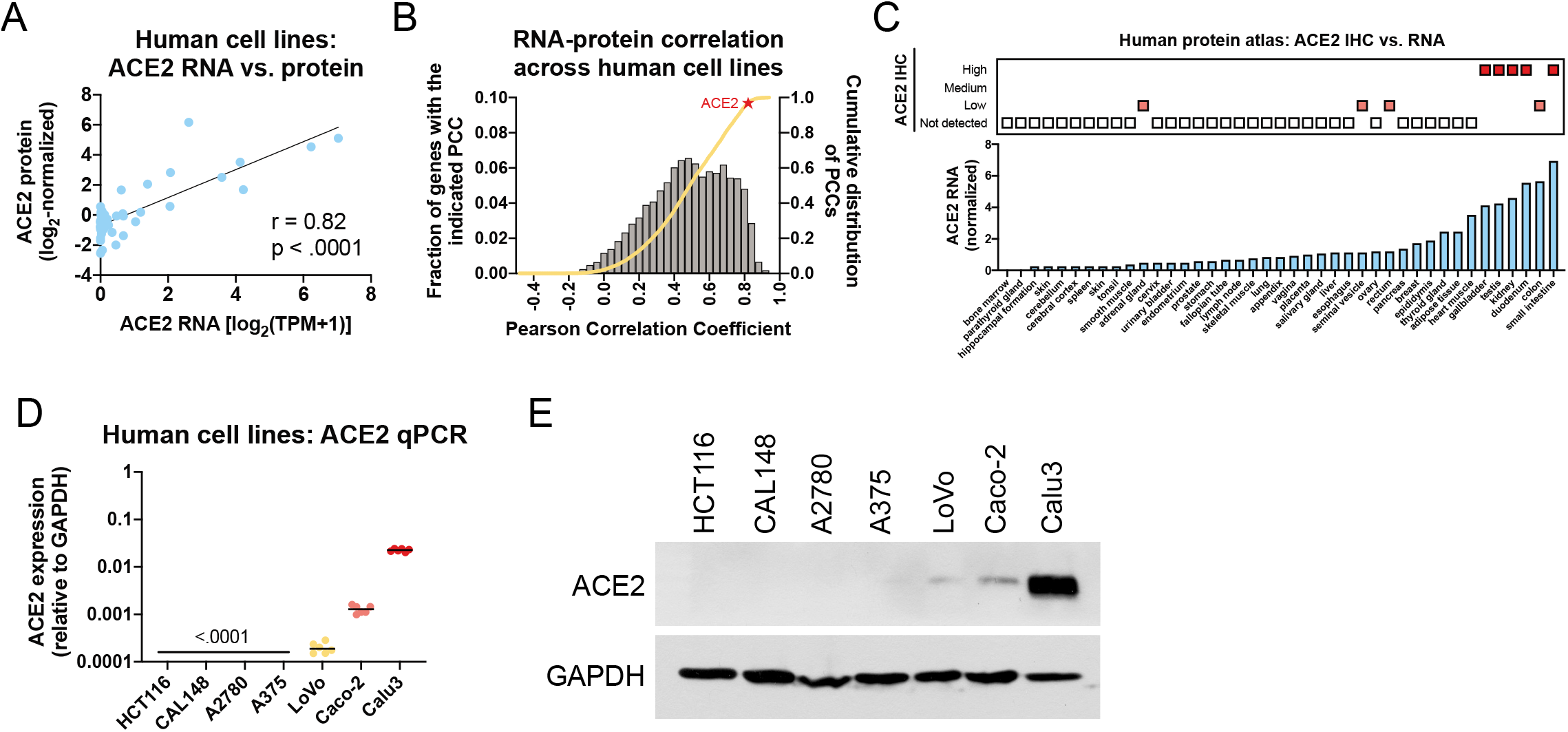
ACE2 RNA levels and protein levels are tightly correlated. Related to Figure 5. (A) A scatter plot comparing the expression of ACE2 RNA, as measured by RNA-Seq, and ACE2 protein, as measured by mass spectrometry, across human cancer cell lines. The black line indicates a linear regression plotted against the data. (B) A bar graph comparing the Pearson correlation coefficients between RNA and protein expression across 11,362 human genes. The grey bars denote the fraction of genes with an indicated correlation coefficient while the orange line indicates the cumulative distribution of those coefficients. ACE2’s position is indicated with a red star. (C) A graph comparing ACE2 immunohistochemistry staining in various tissues vs ACE2 RNA expression in those same tissues. (D) qRT-PCR of ACE2 expression in the indicated cell lines, relative to GAPDH expression. (E) Western blotting of ACE2 expression in the indicated cell lines. GAPDH was analyzed as a loading control. The data in A and B are from ref^106^. The data in C is from the Human Protein Atlas^138^. Additional information on these sources is included in Table S1.

**Table S1. Data sources used for this analysis.**

**Table S2. ACE2-correlated genes and GO terms.**

